# Structural basis of cage assembly by a bacterial flotillin

**DOI:** 10.64898/2026.11.15.766717

**Authors:** Naseer Iqbal, Alireza Ghanbarpour

**Affiliations:** Department of Biochemistry and Molecular Biophysics, Washington University School of Medicine, St. Louis

## Abstract

Flotillins organize membrane microdomains implicated in signaling, cell-wall synthesis, and antibiotic resistance, yet how bacterial flotillins assemble remains unknown. Here, we report a 3.1 Å cryo-EM structure of the *Escherichia coli* flotillin YqiK, purified through its NfeD-family partner YqiJ – the first high-resolution structure of a bacterial flotillin assembly. YqiK forms a membrane assembly of 54 identical subunits. Unlike human flotillin cages, which contain two alternating paralogues, YqiK assembles from a single transmembrane protein. The cage is ~38 nm wide and encloses ~900 nm^2^ of membrane within its ring of 54 transmembrane helices. Its wall helices run continuously for ~287 Å, the longest in any SPFH assembly, making this the largest membrane-anchored SPFH cage reported to date. Helix length and tilt set cage dimensions, while intersubunit salt bridges and hydrophobic interactions stabilize the assembly. Focused classification revealed local discontinuities in the cage wall, suggesting possible conformational flexibility. Mass spectrometry identified stress-associated membrane proteins that co-purify with YqiK–YqiJ. Together, these findings reveal how a bacterial flotillin builds a large membrane-associated scaffold and provide a structural basis for investigating how such compartments organize membrane proteins within the microdomain they define.

## INTRODUCTION

Bacterial membranes are organized into functional membrane microdomains that, like eukaryotic lipid rafts, are scaffolded by flotillins (Lopez & Koch, 2017). These microdomains gather proteins involved in signal transduction, protease secretion and biofilm formation, and they act as platforms on which membrane protein complexes oligomerize (Lopez & Koch, 2017). Their importance is clearest in pathogens. In methicillin-resistant *Staphylococcus aureus* (MRSA), flotillin is required for oligomerization of PBP2a, the enzyme responsible for penicillin resistance, and drugs that disassemble flotillin microdomains make MRSA infections susceptible to penicillin(García-Fernández *et al*, 2017). In *Bacillus subtilis*, flotillins increase membrane fluidity and thereby control the movement of the cell-wall synthesis machinery during fast growth by altering lipid packing behaviour (Zielińska *et al*, 2020). Flotillin micro-domains also act as membrane chaperones: in *S. aureus*, the flotillin FloA and its NfeD partner form a complex whose flexible coiled-coil “ten-tacles” hold unfolded client proteins, including PBP2a, and stabilize them without ATP. Cells lacking FloA or NfeD survive oxidative stress poorly, are attenuated in infection models and lose penicillin resistance(Ukleja *et al*, 2024).

Flotillins belong to the SPFH (stomatin, prohibitin, flotillin, HflK/C) family, which is found in all domains of life (Daumke & Lewin, 2022; Hinderhofer *et al*, 2009). Cryo-EM structures now show a shared architecture in which membrane-associated SPFH domains form the base and extended helical segments assemble with neighbouring subunits into a cup-shaped cage (Gao *et al*, 2025; Lu *et al*, 2026; Ma *et al*, 2022; Qiao *et al*, 2022; Stoner *et al*, 2025). The number of subunits varies widely: 16 in human stomatin (Stoner *et al*., 2025), 22 in the prohibitin PHB1/PHB2 complex (Lange *et al*, 2025) and 26 in the erlin1/2 complex (Gao *et al*., 2025), 24 in *E. coli* HflK/C (Ghanbarpour *et al*, 2025; Iqbal *et al*, 2026; Ma *et al*., 2022; Qiao *et al*., 2022) and 44 in the human flotillin-1/2 complex (Fu & MacKinnon, 2024).

HflK/C is the best-studied bacterial SPFH assembly. HflK and HflC alternate in a 24-subunit assembly in the periplasm that surrounds the membrane-embedded AAA+ protease FtsH (Ma *et al*., 2022; Qiao *et al*., 2022). We previously showed that HflK/C forms not only a closed, symmetric vault but also an asymmetric, nautilus-like shell around one or two FtsH hexamers (Ghanbarpour *et al*., 2025). The shell leaves a ~70-Å passage through which membrane substrates can reach FtsH, and it curves the enclosed membrane, a deformation associated with increased lipid flip-flop (Ghanbarpour *et al*., 2025). We further showed that the membrane-bound FtsH–HflK/C degradation assembly is structurally plastic, and that restricting this flexibility by disulfide-mediated stabilization of the closed conformation impairs bacterial adaptation to aminoglycoside stress (Iqbal *et al*., 2026). Prohibitin has since been shown to adopt an open conformation similar to that of HflK/C (Rose *et al*, 2025). Whether such conformational flexibility extends to bacterial flotillins is unknown.

Structural information on bacterial flotillins is limited to low-resolution reconstructions of the *S. aureus* FloA–NfeD complex, in which two FloA–NfeD units form an open, clamp-shaped assembly (Ukleja *et al*., 2024). How bacterial flotillins assemble to organize membrane micro-domains, and how their architecture relates to that of other SPFH proteins remain less understood. To address these questions, we turned our attention to the *E. coli* flotillin homologue YqiK, which is encoded next to the NfeD-family membrane protein YqiJ (Hinderhofer *et al*., 2009)(SI Appendix, Fig. S1). The only structure of an *E. coli* SPFH–NfeD pair, QmcA–YbbJ (Tan *et al*, 2024), is of a non-flotillin SPFH protein. QmcA forms a 26-subunit cage, and one YbbJ binds each QmcA outside the wall, with its OB-like domain against the SPFH domain and its transmembrane helices against that of QmcA (Tan *et al*., 2024). YqiK belongs to the FloT/eu-karyotic flotillin branch, sharing ~28% sequence identity with *B. subtilis* FloT and 23% with human flotillin-1 and flotillin-2. It is found in detergent-resistant membrane fractions, forms discrete foci in cells and contributes to swimming motility and stress responses (López & Kolter, 2010; Padilla-Vaca *et al*, 2019) including signalling through the Psp membrane-stress response (Jovanovic *et al*, 2014).

Here, we report the first high-resolution cryo-EM structure of YqiK purified through its FLAG-tagged NfeD-family partner YqiJ. YqiK forms a 54-subunit cage and unusually long, continuous wall helices. The structure reveals how a single bacterial flotillin builds the largest membrane-anchored SPFH cage described to date and provides a framework for investigating how flotillin assembly organizes membrane microdo-mains and corrals protein complexes within the large compartment it encloses.

## RESULTS

### Cryo-EM reveals cup-shaped particles following co-expression of YqiK and YqiJ

To test whether bacterial flotillins form a cage-like complex, we first expressed YqiK alone, which formed only small assemblies in negative-stain and cryo-EM screening (SI Appendix, Fig. S2). We therefore turned to YqiJ, which is encoded immediately upstream of *yqiK* in the same operon and belongs to the NfeD family, like YbbJ, the partner of the *E. coli* SPFH protein QmcA (Tan *et al*., 2024). We co-expressed His-tagged YqiK and FLAG-tagged YqiJ from a dual-expression plasmid, and FLAG-YqiJ pulled down YqiK in (n-dodecyl-β-D-maltoside) DDM (SI Appendix, Fig. S1C).

Cryo-EM of the eluate revealed large, cup-shaped particles predominantly in thicker ice, possibly because of their massive size. Refinement with C_27_ symmetry yielded a final reconstruction at a resolution of 3.06 Å, as determined using the gold-standard FSC threshold of 0.143 (Fig. 1; SI Appendix, Fig. S3). The clear resolution of the coiled-coil helices and SPFH domains supported the choice of symmetry, whereas alternative symmetries failed to resolve these features. To improve local resolution in different regions of the map, we subtracted the signal from the remainder of the particle and independently refined the SPFH belt and linker with C_54_ symmetry and the roof with C_27_ symmetry, achieving resolutions of 3.7 and 2.9 Å, respectively (SI Appendix, Figs. S3–S7).

**Fig. 1.**
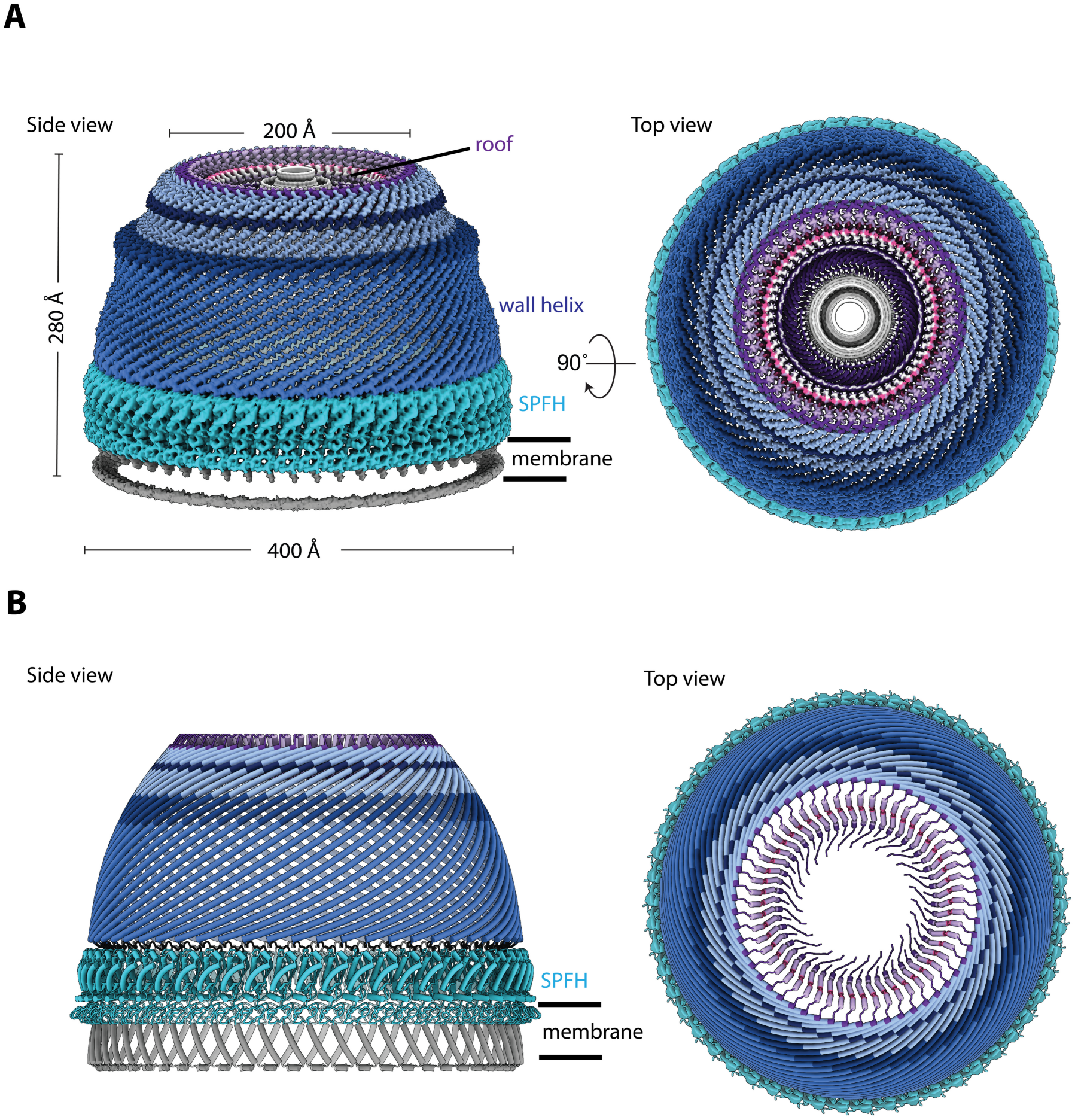
YqiK builds a 54-subunit cage on the cytoplasmic face of the inner membrane. (**A**) Unsharpened cryo-EM density of the YqiK cage, a composite of the 3.1 Å consensus C_27_ map and the SPFH-belt- and roof-focused maps, seen from the side (left) and, after a 90° rotation, down the cage axis from the roof (right). The density is colored by the fitted model: SPFH domains (cyan), wall helices (blue), roof helices and kinks (light purple) and, in the top view, the unassigned density at the center of the roof (grey). The grey ring at the open rim is the detergent micelle and marks the membrane. The cage is ~380 Å across at its widest point, the SPFH belt, and ~278 Å tall. (**B**) The atomic model in the same two views and at the same scale, with the 54 N-terminal transmembrane helices in grey below the SPFH belt.

Representative density is shown in SI Appendix, Fig. S8. The final model, built with ModelAngelo (Jamali *et al*, 2024) into these maps and refined against a composite of the three, covers residues 2–481 (Fig. 2A, Table S1).

**Fig. 2.**
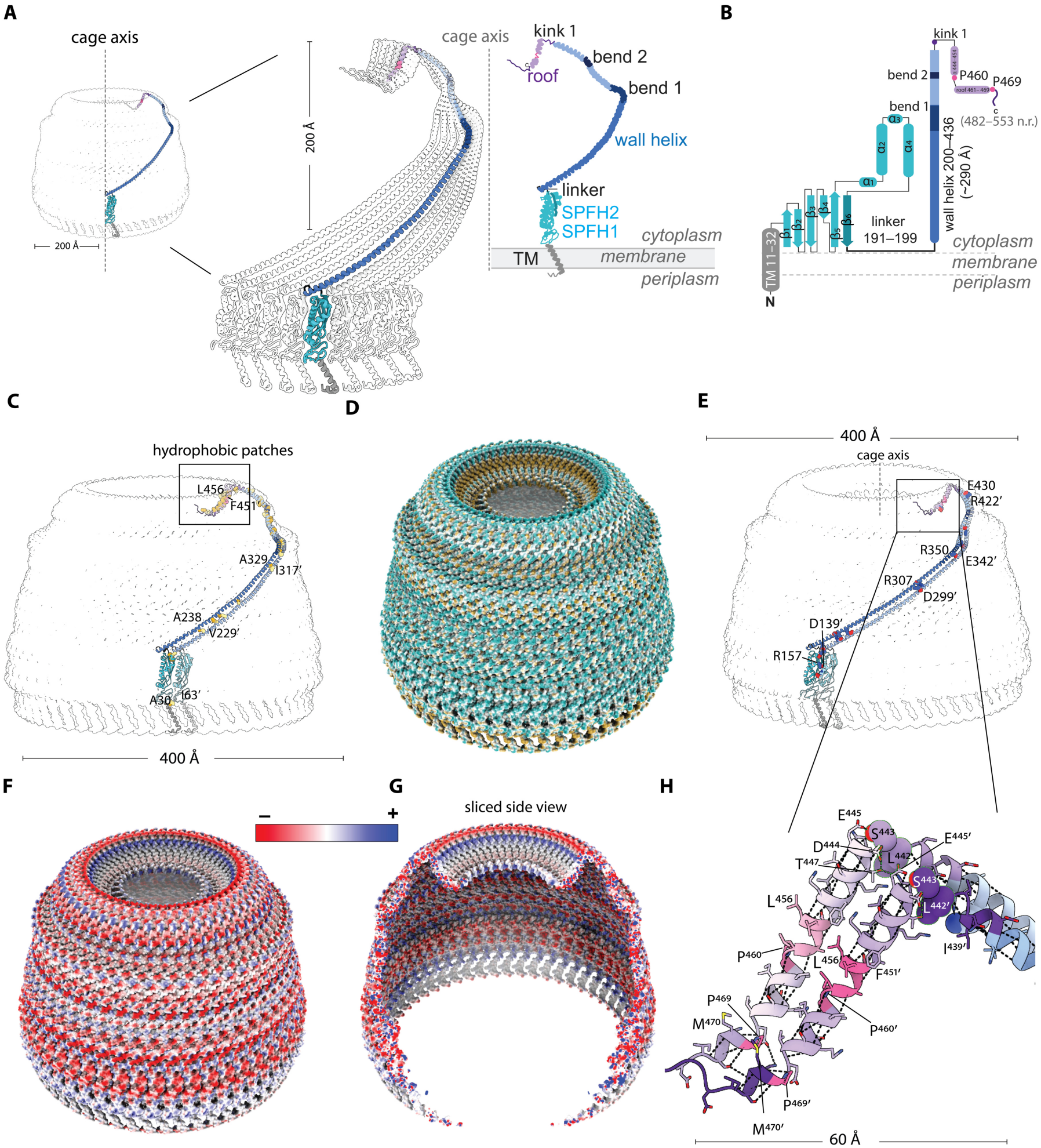
Each YqiK spans the cage wall as a single helix and contacts its two neighbors. (**A**) One YqiK subunit within the cage outline (left), enlarged with surrounding subunits shown as outlines (center), and oriented relative to the membrane (right). The chain is colored by segment: transmembrane helix (TM, grey), SPFH domain (SPFH1 and SPFH2, cyan), wall helix (blue), bends 1 and 2 (dark blue), roof helix (purple) and C-terminal segment (pink). The linker and kink 1 are indicated. (**B**) Topology diagram showing the secondary structure assigned from the model. Residues 482–553 were not resolved (n.r.). (**C**) Inter-subunit hydrophobic contacts along one interface, shown on two neighboring subunits within the cage outline. Contacting side chains are shown as gold spheres. Representative residue pairs are labelled; primes denote the neighboring subunit. (**D**) Molecular surface colored by hydrophobicity, from hydrophilic (cyan) to hydrophobic (gold), highlighting hydrophobic patches across the cage and a hydrophobic ring around the central roof opening. (**E**) Inter-subunit salt bridges along the same interface and in the same orientation as in (**C**). Side chains are shown as spheres (Asp/Glu, red; Lys/Arg, blue), and representative residue pairs are labelled. (**F, G**) Molecular surface colored by Coulombic electrostatic potential, from negative (red) to positive (blue), shown as an external view (**F**) and a sliced side view exposing the cage interior (**G**). (**H**) Enlarged view of the wall-to-roof transition in two neighboring subunits, corresponding to the boxed region in (**E**). Selected residues around kink 1, the roof helices and the C-terminal segments are labelled.

### YqiK builds a 54-subunit cage, the largest membrane-anchored SPFH cage

The map shows a cup-shaped cage ~38 nm across at its widest point, the SPFH belt just above the membrane, and ~28 nm tall from the transmembrane helices to the roof. The 54 transmembrane helices form a ring ~35 nm in diameter, enclosing ~900 nm^2^ of membrane (Fig. 1A, B; Movie S1). The wall contains 54 regularly spaced YqiK subunits, two in each C_27_ repeat. This makes YqiK the largest membrane-anchored SPFH cage described to date; previously reported cages contain 16–44 subunits (Fu & MacKinnon, 2024; Gao *et al*., 2025; Ghanbarpour *et al*., 2025; Ma *et al*., 2022; Stoner *et al*., 2025). The soluble vault, built from 78 copies of the SPFH-like major vault protein, is larger but generally is not membrane-anchored (Lövestam & Scheres, 2025). Within each C_27_ repeat, the two YqiK subunits are indistinguishable in the SPFH ring and wall.

The differences we detect are limited to the extent to which their C-terminal tails are ordered inside the roof (see below) and the low-resolution density above the roof. These differences may account both for the symmetry mismatch between the SPFH belt and the roof, and for the improved refinement of the consensus map under C_27_ symmetry.

### YqiJ is not resolved in the final map

Although the complex was purified through FLAG-tagged YqiJ, YqiJ itself was not resolved in the density. Focused 3D classification of symmetry-expanded particles (*SI Appendix*, Fig. S3) revealed additional density on the outer face of the SPFH belt in 9 of 12 classes, positioned at a different angle around the repeat in each class; additional density was also visible in low-pass-filtered maps of the SPFH belt. Its volume is consistent with the cytoplasmic domain of YqiJ (residues ~123– 209) (*SI Appendix*, Fig. S9). The failure to resolve YqiJ likely reflects several factors acting together: the cytoplasmic OB-fold domain is small (~10 kDa) and contributes little to the alignment signal, it is not held at a fixed position around the repeat, and it may be present at less than one copy per YqiK – so that imposing symmetry averages it out. Consistent with this tentative assignment, AlphaFold3 models of YqiK with YqiJ place the YqiJ OB-fold domain at this position, against the outer face of the SPFH domain, with the YqiJ transmembrane helices beside those of YqiK, much as YbbJ binds QmcA (Tan *et al*., 2024) (*SI Appendix*, Fig. S10). Although model confidence is moderate (ipTM ~0.55), models with one YqiJ per YqiK fit without clashes and are compatible with the cryo-EM data.

### Each YqiK spans the wall as a single helix

At the membrane end, the SPFH domains form a deep belt with approximately 20 Å of circumference per subunit (Fig. 1A, B; Fig. 2A). Each YqiK contains a single N-terminal transmembrane helix (residues ~11–32), preceded by a short acidic sequence (MDD) and flanked at its cytoplasmic end by basic residues (R31, R34, R35). According to the positive-inside rule, this charge distribution predicts a periplasmic N-ter-minus and a cytoplasmic C-terminal region. Consistent with this prediction, a proteome-wide PhoA/GFP fusion analysis of *E. coli* inner membrane proteins experimentally localized the YqiK C-terminus to the cytoplasm (Daley *et al*, 2005). The SPFH domain, wall, and roof therefore face the cytoplasm, as in other flotillins (Fig. 1A; Fig. 2A, B). By comparison, the HflK/C cage extends into the periplasm, illustrating how SPFH proteins assemble cages on either side of the bacterial inner membrane (Ma *et al*., 2022; Qiao *et al*., 2022).

Each SPFH domain (residues 33–190) ends in a β-strand (residues 180–190) that pairs antiparallel with the strand formed by residues 94–104, through backbone hydrogen bonds (Fig. 2B). The focused map of the SPFH belt resolves a nine-residue linker (residues 191–199) as continuous density, so the chain can be traced without a break from each SPFH domain into the base of the wall helix directly above it. This establishes the connectivity between the two regions, which is otherwise ambiguous because 54 SPFH domains and 54 wall helices are inter-leaved around the ring (Fig. 2A, B; SI Appendix, Fig. S8). Above the belt, as in other SPFH assemblies (Gao *et al*., 2025; Lu *et al*., 2026; Ma *et al*., 2022; Qiao *et al*., 2022; Stoner *et al*., 2025), the wall is a single layer of α-helices, one per subunit, with neighboring helices tilted ~58° from the cage axis (Fig. 1B; Fig. 2A). Each helix (residues 200–436) is continuous over ~280 Å, the longest wall helix described in an SPFH cage. Between residues ~337 and ~368 it bends gently (bend 1, Fig. 2A, B), with the i→i+4 hydrogen bonds lengthening at residues 338–340 but the helix continuing without interruption, dividing the wall into a lower (residues 200–~337) and an upper (~368–436) segment; this bend sits where HflK/C, erlin and prohibitin bend between CC1 and CC2 (Gao *et al*., 2025). Even its lower segment (residues 200–337; 138 residues; ~198 Å) exceeds the full wall-helix length of every SPFH assembly except human flotillin-1. A second gentle bend at residues ~401–409 (bend 2, Fig. 2A, B), also with its hydrogen bonds largely intact, turns the upper wall inward toward the kink.

### Salt bridges and hydrophobic interactions hold neighboring subunits together

Every YqiK contacts only the two adjacent subunits, but does so along its full length; each of these two interfaces buries ~8,000 Å^2^ on each subunit. About two-thirds of this interface lies in the wall. Adjacent wall helices are offset by about two helical turns and are joined by a hydrophobic seam, which forms a continuous apolar surface running up the wall to a ring around the central roof opening (Fig. 2C, D), and by a ladder of interhelical salt bridges between residue i of one helix and residues i+8 to i+12 of its neighbor (for example D299–R307, E342–R350, E366– R376 and R422–E430; Fig. 2E). On the molecular surface the lumenal and outer faces of the wall are acidic, so these complementary charges lie in the buried seam rather than on the exposed faces (Fig. 2F, G).

At the top of the wall, residues 437–443 leave the helix: the i→i+4 backbone hydrogen bonds break, L442 and S443 adopt extended conformations, and the chain turns sharply into a second helix that starts at D444 and runs back down along the inside of the wall top, ~90 Å from the cage axis (Fig. 2H). This roof helix is bent by P460 but not interrupted by it, so the chain turns inward and residues 460–469 converge over the cage to form the roof (Fig. 1A, B, top views; Fig. 2B, H). P469 breaks the helix, and beyond P469 the C-terminal segment leaves the roof as extended chain. In the 2.9 Å roof-focused map (SI Appendix, Fig. S6) the C-terminal tails of the two YqiK of each repeat are ordered to different extents: on one, residues 472–481 continue as an extended segment into the roof; on the other, only residues 472–474 are ordered (Fig. 2E, H). Additional ordered density fills the roof ring and fits the remaining C-terminal residues (482– 553), but their sequence could not be confidently assigned, so they were not modelled (Fig. 2B).

### Wall-helix length correlates with SPFH cage size

Comparison of YqiK with other membrane-associated SPFH assemblies reveals a conserved architecture with substantial variation in wall length and cage size (Fig. 3). YqiK and human flotillin-1/2 form similarly shaped cages but differ in membrane anchoring. YqiK is anchored by a ring of 54 transmembrane helices, whereas human flotillin-1/2 lacks transmembrane helices and associates with the bilayer through its acylated SPFH1 domains (Fu & MacKinnon, 2024) (Fig. 3A). Superposition of the SPFH domains gives an RMSD of 1.3 Å over 105 and 83 Cα atoms for flotillin-1 and flotillin-2, respectively, with the wall helices emerging from the domains in similar directions (Fig. 3B).

**Fig. 3.**
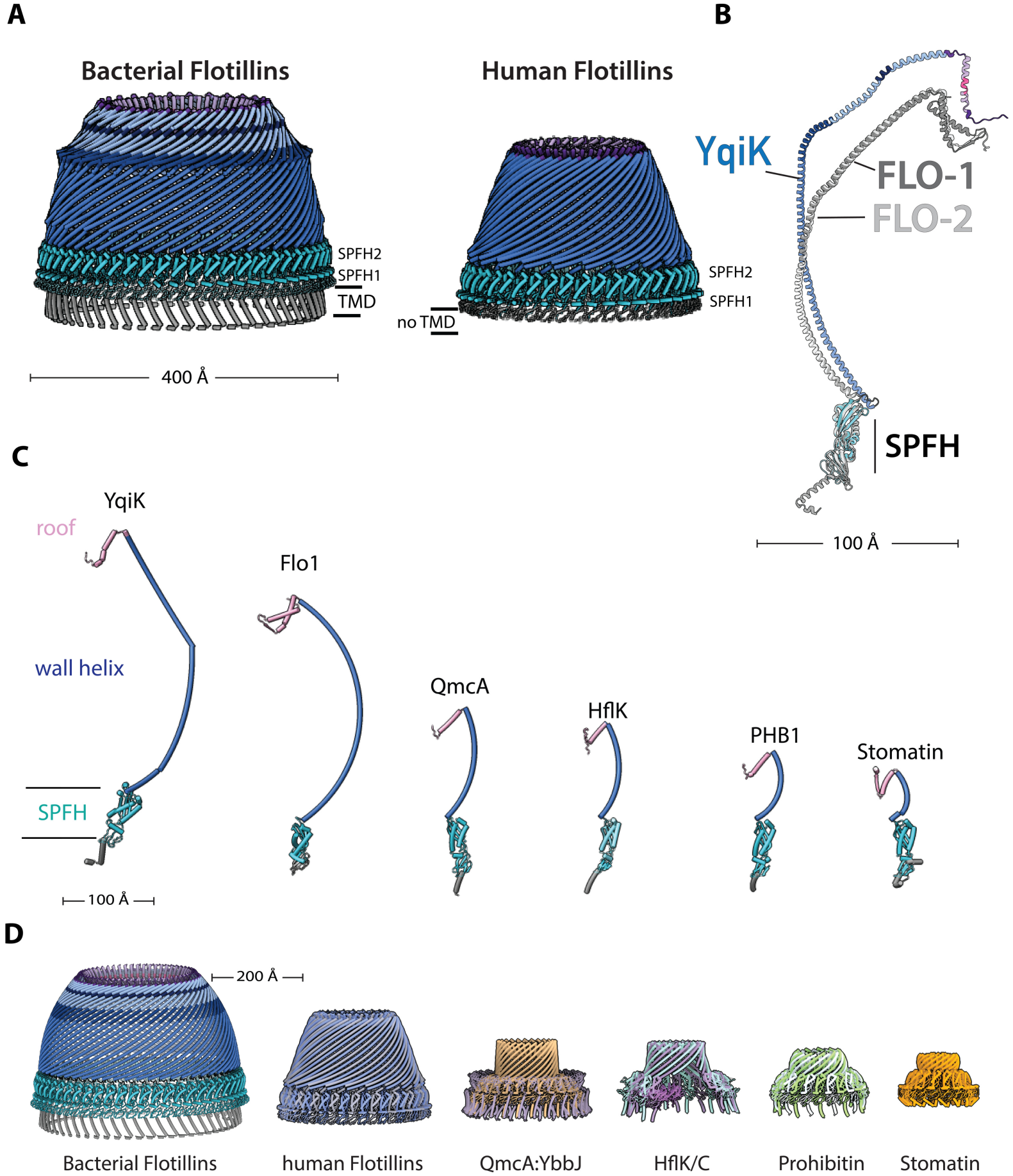
YqiK compared with other membrane-associated SPFH assemblies. (**A**) The YqiK and human flotillin-1/2 cages as cartoons at the same scale, with the SPFH1 and SPFH2 subdomains labelled; YqiK is anchored by a ring of transmembrane helices (TMD, grey) that human flotillin lacks. SPFH ring cyan, wall and roof blue and pink respectively. (**B**) One YqiK subunit (blue) superposed through its SPFH domain on flotillin-1 (dark grey) and flotillin-2 (light grey) of PDB 9L3G. (**C**) One subunit from each assembly, superposed on the SPFH domain and shown at the same scale: YqiK, flotillin-1 (Flo1), QmcA, HflK, prohibitin-1 (PHB1) and stomatin. Colored by region: membrane anchor grey, SPFH domain cyan, wall helix blue, roof and C-terminal segment pink. (**D**) The corresponding SPFH oligomeric assemblies at the same scale, each in a single color. YqiK has the longest wall helix and the largest cage of those shown.

Across the family, the conserved SPFH domain supports helical regions of markedly different lengths (Fig. 3C). Measured end to end, the wall helix spans ~287 Å in YqiK (residues 200–436), ~236 Å in flotillin-1, ~118 Å in QmcA, ~84 Å in HflK, ~76 Å in prohibitin-1 and ~55 Å in stomatin. The corresponding assemblies contain 54, 44, 26, 24, 22 and 16 subunits, respectively (Fig. 3D). This trend is consistent with the reported linear relationship between subunit number and the length of the first coiled-coil segment, CC1, across bacterial and human SPFH complexes (Gao *et al*., 2025). YqiK extends this comparison with a continuous 237-residue wall helix rather than separate CC1 and CC2 helices. Applying the published fit (Gao *et al*., 2025) to the full YqiK wall helix predicts ~53 subunits, close to the 54 observed.

### YqiJ co-purifies with YqiK and with stress-associated membrane systems

Flotillin complexes have been implicated in membrane organization and the assembly of multisubunit protein complexes (Lopez & Koch, 2017). To identify proteins associated with the purified complex, we compared FLAG-YqiJ eluates with empty-vector controls by data-independent-acquisition mass spectrometry (three biological replicates each; 2,212 proteins quantified, 1,873 of them in at least two replicates of both conditions; Dataset S1). YqiJ was the most strongly enriched protein (log2FC 8.7), and YqiK was recovered with it (log2FC 5.8), as expected for the purified complex (Fig. 4A; *SI Appendix*, Fig. S1). The proteins enriched alongside them were components of membrane systems that *E. coli* deploys under stress. ElaB and YqjD are tail-anchored DUF883 proteins that accumulate in stationary phase and localize to the inner membrane (Guo *et al*, 2017; Yoshida *et al*, 2012).

**Fig. 4.**
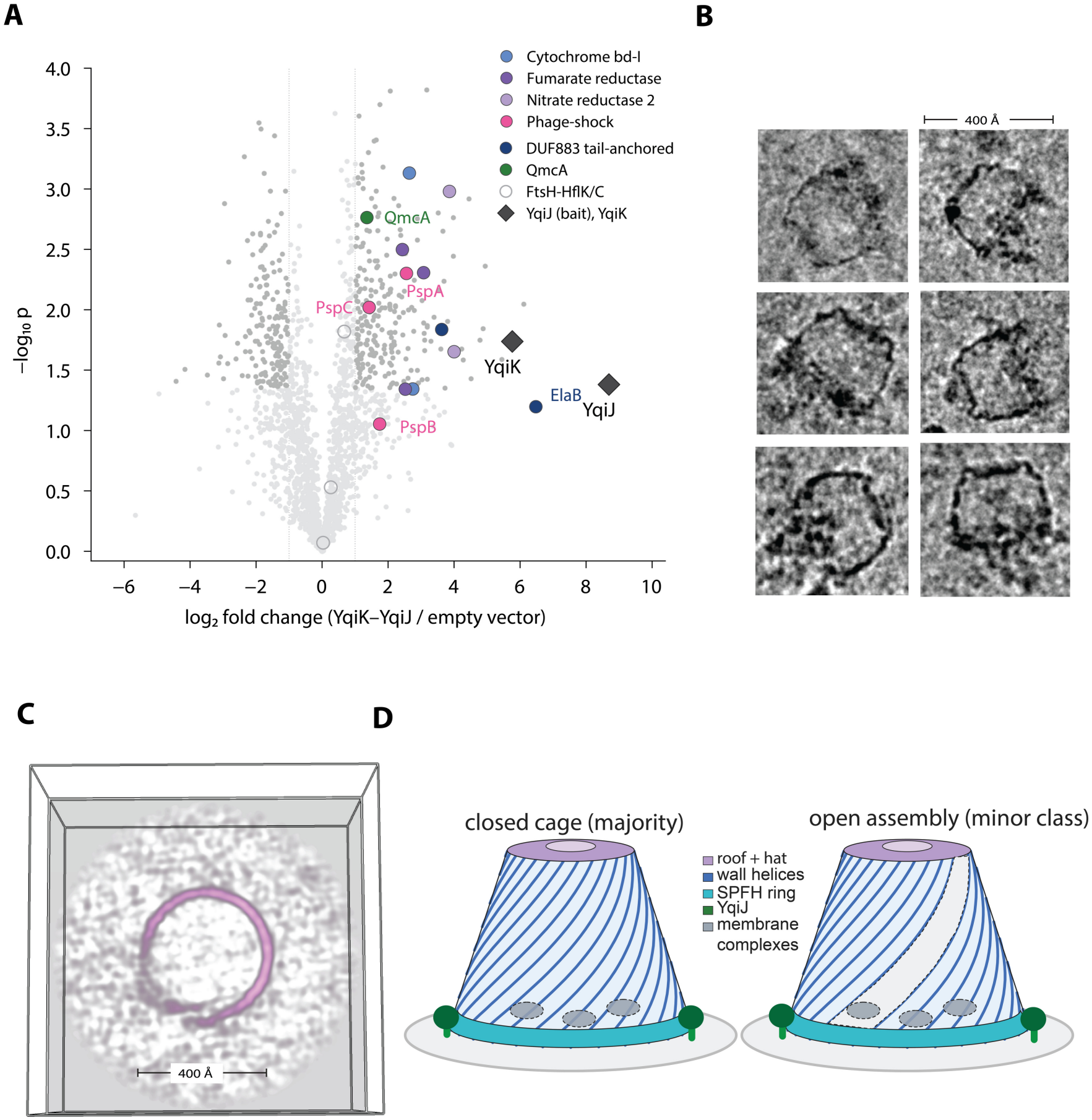
YqiK cages enclose additional material and co-purify with stress-related membrane systems. (**A**) Volcano plot of FLAG-YqiJ eluates against empty-vector controls (three biological replicates). Colored circles mark the complexes in the key; YqiJ and YqiK are labelled. Axes are truncated for clarity: nine protein groups with −log_10_ p > 4 lie above the plotted range. All tested protein groups are listed in Dataset S1. (**B**) Representative cryo-EM particle images containing additional density projected within the cage outline. (**C**) A slab through the cage, at a z position chosen to cut the wall, of a representative class from 3D classification without alignment (SI Appendix, Fig. S11) showing an open conformation. Wall density is absent along one intersubunit seam while the rest of the ring stays continuous. The outline marks the box size. (**D**) Schematic illustrating a proposed model in which cage flexibility could allow membrane proteins to enter and become concentrated within the enclosed membrane region. The closed cage and putative open state are colored as indicated in the key.

NarY and NarZ are subunits of nitrate reductase Z, and FrdA and FrdC of fumarate reductase, the terminal reductases of anaerobic respiration (Unden & Bongaerts, 1997); CydA and CydB form cytochrome *bd*-I, the high-affinity terminal oxidase induced under microaerobic and stress conditions (Giuffrè *et al*, 2014). PspA and PspC belong to the phage-shock response which is triggered by loss of inner-membrane integrity (Flores-Kim & Darwin, 2016). RpoS was itself enriched, so part of this signature likely reflects stress induction in the expressing cells (Chang *et al*, 1999). By comparison, abundant housekeeping membrane complexes, including NADH de-hydrogenase I and the Sec translocon, showed only modest shifts, as did FtsH and its SPFH partners HflK and HflC. The SPFH paralogue QmcA was modestly enriched. This pattern argues against indiscriminate retention of abundant inner-membrane proteins by the cage.

Visual inspection of individual particles in cryo-EM micrographs revealed additional density at the membrane-proximal end of the cage and within its lumen (Fig. 4B). These features raise the possibility that some of the complexes detected by mass spectrometry remain associated with the assembly. However, the additional density was not resolved in the final reconstruction, possibly because of the imposed symmetry and conformational and compositional heterogeneity of these assemblies. Together, these observations identify proteins that co-purify with the cage and suggest the presence of additional associated material, but whether any of these proteins occupies the membrane domain enclosed by the cage remains to be tested.

### Focused classification reveals local discontinuities in wall density

Previous studies have shown that HflK/C (Ghanbarpour *et al*., 2025) and prohibitin (Rose *et al*., 2025) complexes can adopt open conformations. We therefore asked whether YqiK assemblies also exhibit local openings. We expanded the 5,784 particles using C_27_ symmetry, yielding 156,168 symmetry-related copies, and classified them without alignment into ten classes using a mask focused on one segment of the cage wall (SI Appendix, Fig. S11). Most classes showed continuous wall density. In two classes (3.2% and 4.3% of the expanded set, respectively) density was absent along a band extending from SPFH belt to the roof. A slab through the affected region revealed a gap in the ring of wall helices along a single intersubunit seam (Fig. 4C). The limited resolution of these classes precludes distinguishing separation of neighboring subunits from local disorder or subunit loss. Nevertheless, this discontinuity is consistent with partially open assemblies, analogous to the open conformations reported for HflK/C (Ghanbarpour *et al*., 2025) and prohibitin (Rose *et al*., 2025) complexes.

## DISCUSSION

Bacterial flotillins have long been implicated in membrane microdomain formation, but the organization of their higher-order assemblies has remained unclear. The YqiK cage shares its overall architecture with the human flotillin-1/2 cage (Fu & MacKinnon, 2024; Lu *et al*., 2026), including a membrane-associated SPFH ring, an extended helical wall and a roof (Fig. 3). The human cage contains 44 subunits arranged as alternating flotillin-1 and flotillin-2 paralogues (Fu & MacKinnon, 2024; Lu *et al*., 2026). The YqiK cage contains 54 copies of a single flotillin. Its unusually long wall helices and large subunit count are consistent with the association between helix length and cage size observed across SPFH assemblies (Gao *et al*., 2025; Lu *et al*., 2026; Ma *et al*., 2022; Qiao *et al*., 2022; Stoner *et al*., 2025). NfeD-family partners can contribute to flotillin assembly and function (Ukleja et al., 2024). We recovered large YqiK cages after co-expression with YqiJ and observed peripheral density consistent with the predicted placement of the YqiJ OB-fold domain outside the SPFH belt (SI Appendix, Fig. S9). In this model, YqiJ sits near the membrane, where it could support cage assembly or interact with other proteins. The ability of the *S. aureus* FloA–NfeD complex to stabilize unfolded client proteins suggests that NfeD-family partners may link the organization of flotillin microdomains to protein quality control (Ukleja *et al*., 2024).

What does a cage this large enclose? Its membrane footprint raises the possibility that it concentrates membrane proteins and promotes the assembly or stabilization of multisubunit complexes. Our proteomic analysis detected components of the phage-shock response and respiratory systems in FLAG–YqiJ eluates at higher abundance than in empty-vector controls. FtsH and its SPFH partners HflK and HflC showed smaller changes in abundance. The recovery of phage-shock proteins is consistent with the previously reported role of YqiK in Psp signaling (Jovanovic *et al*., 2014). These proteins are candidates for testing whether they associate with YqiK cages in cells and whether that association affects their activity.

The function of flotillin cages may also depend on their structural flexibility. In a subset of YqiK classes, local gaps in wall density are consistent with separation of neighboring subunits, local disorder or subunit loss (Fig. 4C). If the cage can open reversibly, a gap could allow membrane proteins to move between the region bounded by the cage and the surrounding membrane (Fig. 4D). The open structures of HflK/C and prohibitin motivate testing whether YqiK can undergo a similar transition.

Finally, the YqiK structure provides a template for locating flotillin cages in their native cellular environment using cryo-ET. Identifying these compartments in cells and defining their composition will help connect cage architecture to physiological function. This first near-atomic structure of a bacterial flotillin cage provides a molecular basis for investigating how related scaffolds assemble in other prokaryotes and how membrane organization contributes to cellular adaptation to stress.

### Materials and Methods

#### Plasmid construction

*yqiK* and *yqiJ* were amplified from *E. coli* MG1655 genomic DNA and assembled into pETDuet by Gibson assembly, with an N-terminal His tag on YqiK and a C-terminal FLAG tag on YqiJ.

#### Expression and purification

*E. coli* BL21(DE3) carrying the plasmid was grown in LB with 50 µg/mL ampicillin at 37 °C to an OD_600_ of 0.7, induced with 0.3 mM IPTG and grown for 18 hours at 18 °C. Cells were resuspended in 50 mM Tris-HCl pH 8.0, 100 mM KCl, 10% glycerol (buffer A), flash-frozen, thawed, supplemented with 3 mM β-mercaptoethanol and 5 mM MgCl_2_, and lysed by sonication. After removal of debris (JA-20 rotor, 14,000 rpm, 30 min, 4 °C), membranes were pelleted (Ti45 rotor, 30,000 rpm, 1 hour, 4 °C), resuspended in buffer A with 5 mM MgCl_2_ and 3 mM β-mercaptoethanol, Dounce-homogenized and solubilized in 1% DDM for 2 hours at 4 °C. After ultracentrifugation (Ti45, 25,000 rpm, 45 min), the supernatant was incubated with anti-FLAG M2 resin (Millipore Sigma A2220) for 2 hours at 4 °C, washed with 3 mL of the same buffer containing 0.03% DDM, and eluted with 0.25 mg/mL FLAG peptide (APExBIO) in that buffer. Eluates were concentrated on a 100-kDa cutoff centrifugal concentrator (Millipore).

Protein identity was confirmed by anti-His and anti-FLAG immunoblotting and by mass spectrometry.

#### Proteomics

FLAG-YqiJ eluates and eluates from cells carrying the empty vector, prepared identically (three biological replicates each), and analyzed by data-independent-acquisition LC-MS/MS at the Mass Spectrometry Technology Access Center at the McDonnell Genome Institute (MTAC@MGI), Washington University School of Medicine.

#### On-bead digestion

Trypsin (800 ng) was added to immunoprecipitated proteins on the beads for 6 hours at 37 °C. Digested samples were centrifuged for 2 min at 5,000 × g and the supernatants collected. Beads were washed twice with 100 mM ammonium bicarbonate and the supernatants pooled. Samples were reduced with 20 mM dithiothreitol at 37 °C for 1 hour and cysteines alkylated with 80 mM iodoacetamide for 45 min in the dark, then treated with 600 ng trypsin and incubated overnight at 37 °C. Peptides were desalted by solid-phase extraction on a C18 spin column (Pierce) and eluted with 0.1% formic acid in 80% acetonitrile.

### LC-MS and data analysis

Samples were resuspended in 20 µl of 0.1% Formic acid (v/v) in water, and 5 µl was analyzed using a timsTOF Pro2 Mass Spectrometer (Bruker Daltonics) coupled to a nanoElute2 (Bruker Daltonics) LC system. Peptides were separated on a PepSep C18 column (25 cm length, 150 μm I.D., 1.5 μm particle size, 100 Å pore size; Bruker Daltonics), using a 3% to 35% gradient of Solvent B over 54 min, with a total run time of 60 minutes at a 600 nL/min flow rate. The raw data were collected in the m/z range 100-1700 m/z and mobility range (1/K0) = 0.65-1.38 Vs cm−2 in the dia-PASEF mode. Briefly, the parameters were: number of MS1 ramps = 1, number of MS/MS ramps = 14, number of MS/MS windows = 28. The estimated cycle time was 1.20 s. Both the ramp and accumulation times were set to 75 ms. Raw files were processed using Spectronaut (ver. 21.0) and searched against the UniProt *E*.*coli K12* protein database. Search parameters included Trypsin/P specificity with up to 2 missed cleavages. Carbamidomethylation of cysteine was specified as a fixed modification, and methionine oxidation and N-terminal acetylation were specified as variable modifications. Peptide spectrum matches, peptides, and protein groups were identified using a false discovery rate (FDR) threshold of 0.01.

Protein quantities were calculated from the precursor quantities, calculated as the area under the curve between the XIC peak boundaries for the target ion at the MS2 level, and label-free quantification was performed using Spectronaut (ver. 21.0) with the default directDIA+ (Deep) library-free workflow and searched against the UniProt *E. coli* K-12 protein database (downloaded October 2026).

#### Differential analysis

Differential abundance between FLAG-YqiJ and empty-vector eluates was determined in Spectronaut using its default candidates workflow, which reports the average log2 ratio and p-value for each protein group quantified in at least two replicates of both conditions (Dataset S1).

#### Negative-staining

YqiK samples were prepared for negative-stain electron microscopy using 200-mesh copper grids coated with carbon film (01840-F, Ted Pella.) Grids were glow-discharged for 30s in a Solarus 950 plasma cleaner (Gatan, Pleasanton, CA). A 10 μL aliquot of 10-fold diluted YqiK sample was applied to the carbon-coated side of the grid and incubated for 2 min. Subsequently, the grids were washed five times with 50 μL of ultrapure water and then stained with 50 μL of 0.75% (w/v) uranyl formate for 2 min. Excess liquid was removed by blotting with filter paper (Whatman No.2, Fisher Scientific), and the grids were allowed to air dry. Images were acquired using a JEOL JEM-1400 transmission electron microscope at 40,000 x magnification, operated at an accelerating voltage of 120 kV.

#### Cryo-EM sample preparation

Sample (2.5 µL) was applied to 200-mesh Quantifoil R2/1 copper grids glow-discharged at 25 mA for 30 s (PELCO easiGlow), blotted for 4 s at blot force 0, 6 °C and 100% humidity, and plunge-frozen in liquid ethane cooled by liquid nitrogen using a Vitrobot Mark IV (Thermo Fisher Scientific).

#### Cryo-EM Data collection

A total of 18,457 movies were collected on a Titan Krios at 300 kV with a Falcon 4 detector using EPU (six images per hole; 0.93 Å/pixel; 50.12 e^−^/Å^2^; defocus −0.4 to −2.2 µm).

#### Cryo-EM Data Processing

Data were processed in cryoSPARC v5 (Punjani *et al*, 2017) (SI Appendix, Fig. S3-S7). After patch motion correction and patch CTF estimation, particles picked manually from ~429 micrographs were used to train a Topaz model. Of 13,349 Topaz-picked particles (700-px box, binned to 128 px), 8,889 selected by 2D classification served as templates for template picking. After repeated 2D classification, 5,873 particles were re-extracted (1,012-px box, binned to 640 px). An *ab initio* reconstruction with C_26_ symmetry was followed by non-uniform refinement with C_18_ symmetry, local refinement, local CTF refinement and a second local refinement (3.4 Å). Following C_18_ symmetry expansion and 3D classification using a mask focused on the SPFH belt, a class volume showing YqiJ density was selected as the reference for non-uniform refinement of all 5,873 particles with C_27_ symmetry, yielding a map at ~3.2 Å resolution.

Particles were then re-extracted (960-px box, binned to 684 px; 1.3053 Å/pixel), and an *ab initio* reconstruction with C_27_ symmetry was followed by local refinement, local CTF refinement and a final local refinement with a mask generated in UCSF ChimeraX, giving a final map at 3.06 Å (gold-standard FSC 0.143, automatically tightened mask).

#### Focused maps, hand and composite map

For focused refinement of the SPFH belt, density outside the TM helices, SPFH ring, linker and base of the wall (extending to approximately residue 250) was removed by signal subtraction. The signal-subtracted particles were then recentered and locally refined with C_54_ symmetry, yielding a ~3.7 Å map. For the roof, density outside the roof and the region above it was subtracted, and local refinement with C_27_ symmetry yielded a ~2.9 Å map. The cryoSPARC maps had inverted handedness; all maps were therefore mirrored along the x axis before model building. The corrected hand was confirmed by the fit of the model backbone to the density.

#### Further classifications

To detect open cages, the 5,784 particles were expanded with C_27_ symmetry, giving 156,168 symmetry-related copies, which were classified without alignment into ten classes with a mask on one segment of the cage wall (SI Appendix, Fig. S11).

#### Model building

The TM helix and SPFH domain were taken from the AlphaFold model of YqiK and refined in ISOLDE (Croll, 2018) and Phenix (Afonine *et al*, 2018). The wall, kink and roof (residues 203–481) were built with ModelAngelo (v1.0.18)(Jamali *et al*., 2024) into the sharpened consensus map and, independently, into the roof-focused map; the two builds gave the same sequence register across residues 371– 471. SPFH2, the linker and the base of the wall (residues 91–230) were built with ModelAngelo into the SPFH-focused map. In each case, one ModelAngelo chain close to the consensus of all copies was placed on every subunit with the cage symmetry; residues 472–481 were built on alternate subunits and 472–474 on the others. The final model (residues 2–481 or 2–474) was refined against the composite map in phe-nix.real_space_refine with NCS constraints and secondary-structure and Ramachandran restraints (Table S1). Figures were prepared with UCSF ChimeraX (Meng *et al*, 2023).

## Supporting information

Movie S1. Overall structure of the YqiK-YqiJ cage.

## Data availability

The maps have been deposited in the Electron Microscopy Data Bank with their half maps and masks: consensus C_27_ map, EMD-79930; SPFH-focused map (SPFH belt and transmembrane ring), EMD-79931; roof-focused map, EMD-79939; composite map, EMD-79940. The atomic model has been deposited in the Protein Data Bank (PDB 40AZ; extended ID pdb_000040az) together with the composite map, and the mass spectrometry proteomics data in the ProteomeXchange Consortium via the MassIVE partner repository (PXD087383; MassIVE MSV000104018). Protein quantification and statistics are in Dataset S1.

## COI

The authors declare no conflicts of interest.

## Acknowledgments

This work was supported by startup funds from the Department of Bio-chemistry and Molecular Biophysics at Washington University School of Medicine, and by NIH grant R35-GM160328. Cryo-EM data were collected at the Electron Microscopy Core at the University of Missouri, which is supported by the Office of Research, Innovation, and Impact at the University of Missouri. The authors also thank Han-I Yeh from the Electron Microscopy Core for their technical support. Negative staining data were collected at the Washington University in St. Louis Center for Cellular Imaging (WUCCI). Mass Spectrometry analyses were performed by the Mass Spectrometry Technology Access Center at the McDonnell Genome Institute (MTAC@MGI) at Washington University School of Medicine, supported by the Di-abetes Research Center/NIH grant P30 DK020579, Institute of Clinical and Translational Sciences/NCATS CTSA award UL1 TR002345, and Siteman Cancer Center/NCI CCSG grant P30 CA091842. We thank SBGrid (Morin *et al*, 2013) for providing the structural biology software packages used in this study.

## Author Contributions

N.I. expressed and purified the proteins, prepared the cryo-EM grids. N.I. and A.G. processed the EM data. A.G. performed the final reconstruction and refinement, built and refined the structural models, and wrote the first draft of the manuscript. All authors contributed to writing and editing the manuscript. A.G. supervised the project.

## Supporting Information

**Supplementary Figure 1.**
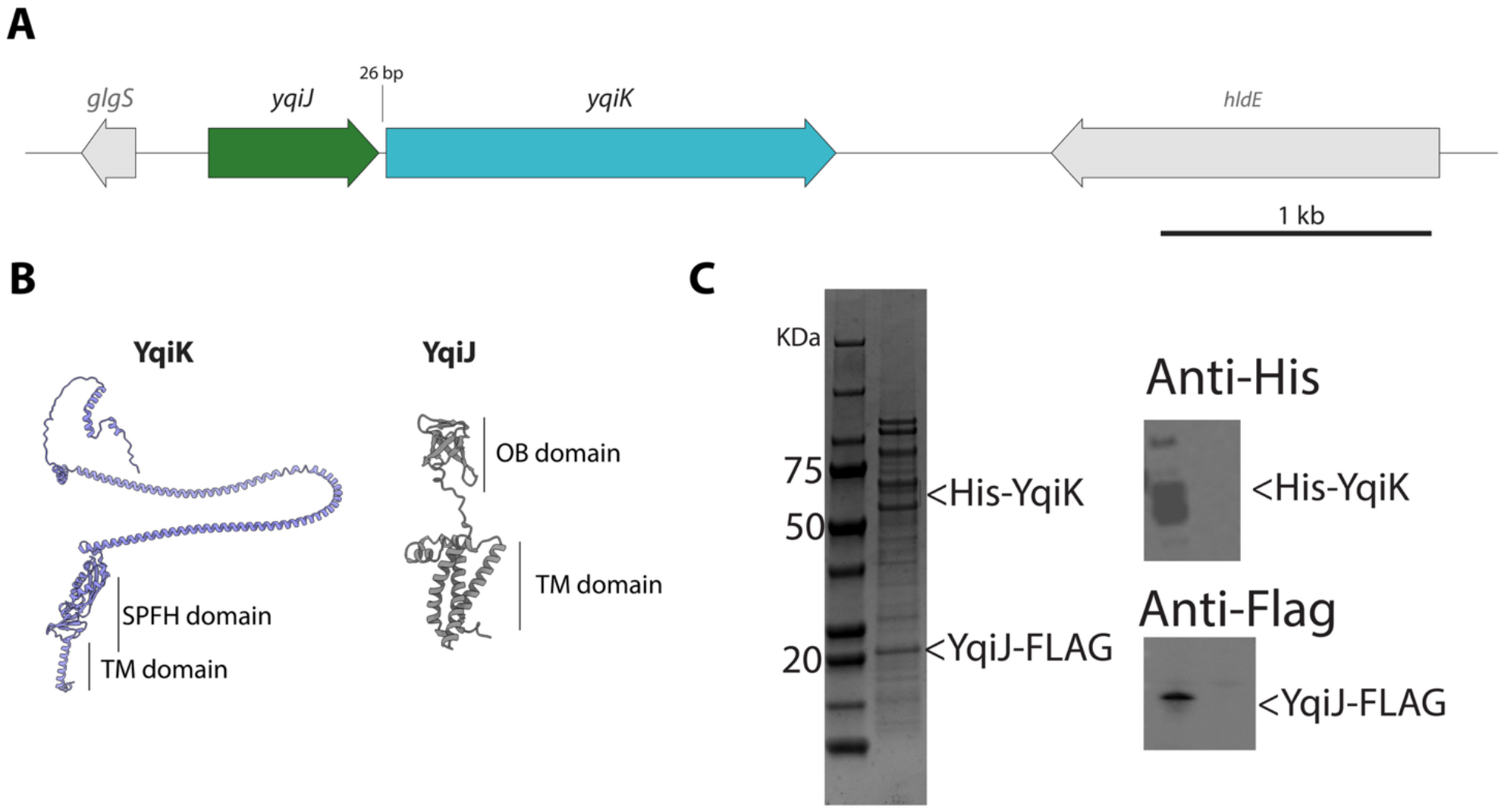
Genomic organization, predicted structures and co-purification of YqiK and YqiJ. (**A**) Genomic organization of the *E. coli yqiJ–yqiK* locus. Arrows indicate gene orientation; *yqiJ* and *yqiK* are separated by 26 bp. (**B**) AlphaFold 3 predicted structures of YqiK and YqiJ, showing the SPFH and transmembrane (TM) domains of YqiK and the OB-fold and TM domains of YqiJ. (**C**) SDS–PAGE analysis and anti-His and anti-FLAG immunoblots of the FLAG-affinity eluate from cells co-expressing His-tagged YqiK and FLAG-tagged YqiJ. Molecular masses are indicated in kDa.

**Supplementary Figure 2.**
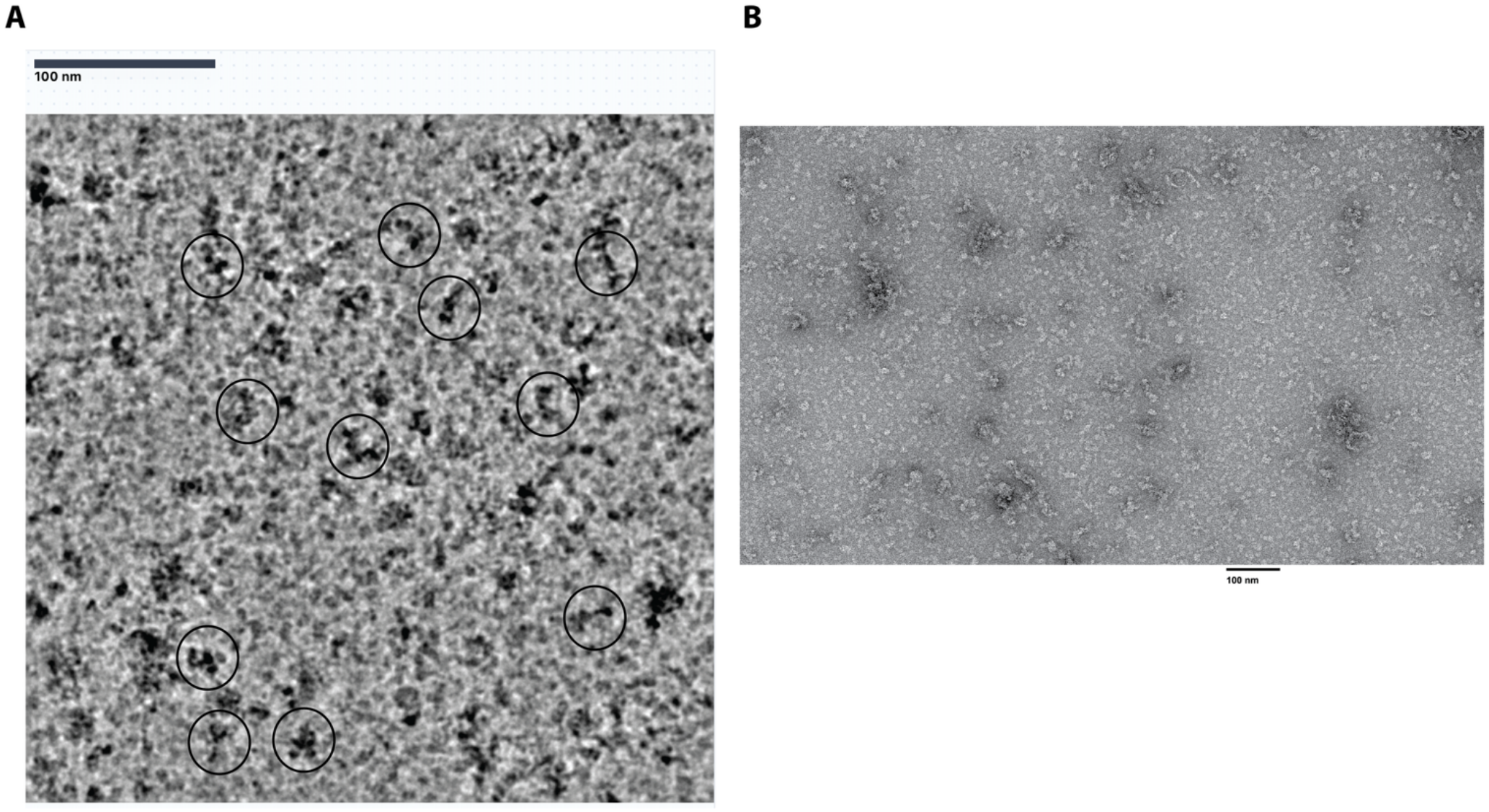
Cryo-EM and negative-stain EM screening of YqiK expressed alone. (**A**) Representative cryo-EM micrograph collected using a Glacios microscope. Circles highlight selected particles. (**B**) Representative negative-stain EM micrograph of the purified sample. Screening revealed small assemblies without the large, cup-shaped cages observed following co-expression with YqiJ.

**Supplementary Figure 3.**
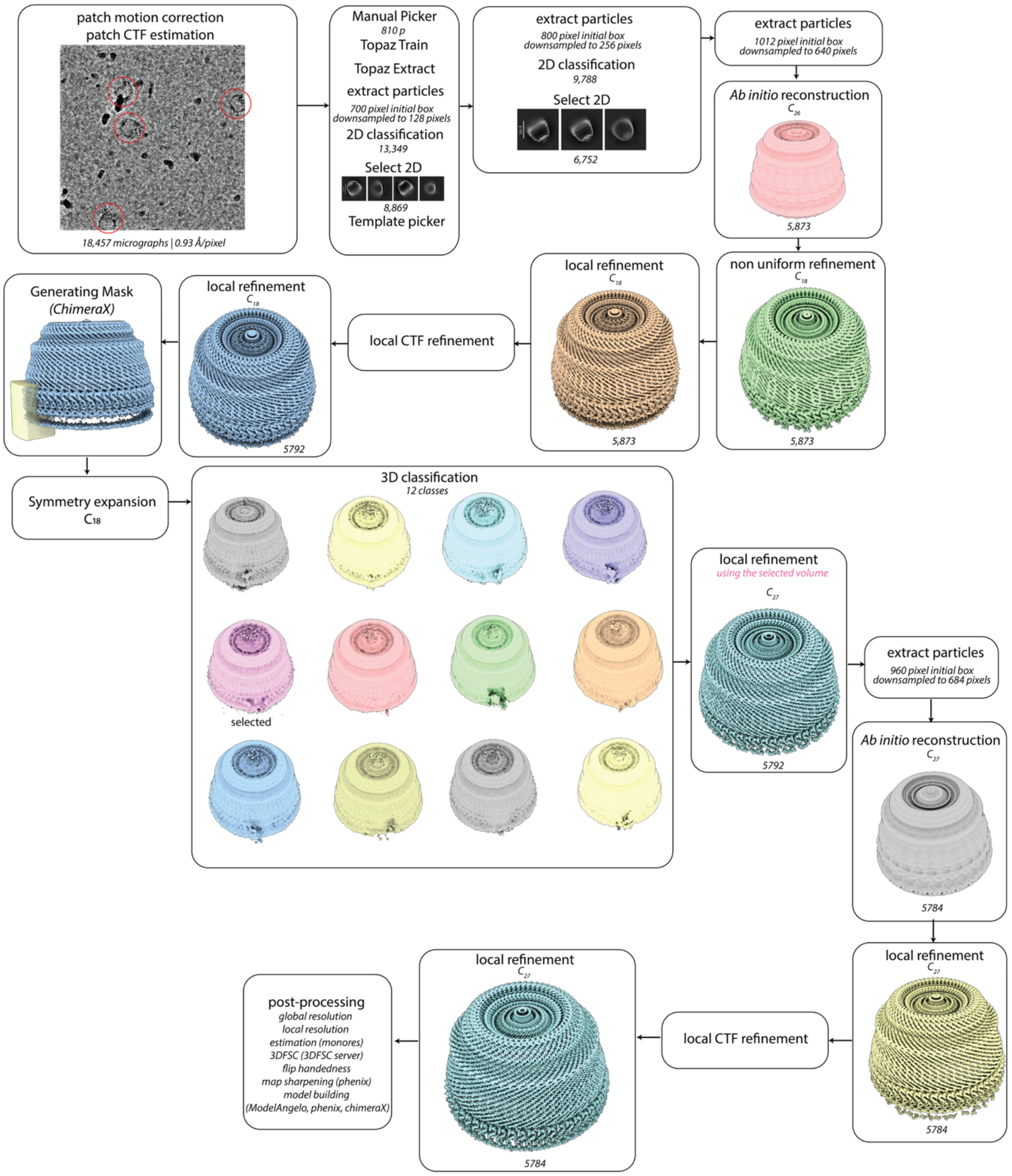
Cryo-EM data-processing workflow for the full YqiK cage reconstruction. Job names, processing details and non-default parameters (italicized) are indicated in each box.

**Supplementary Figure 4.**
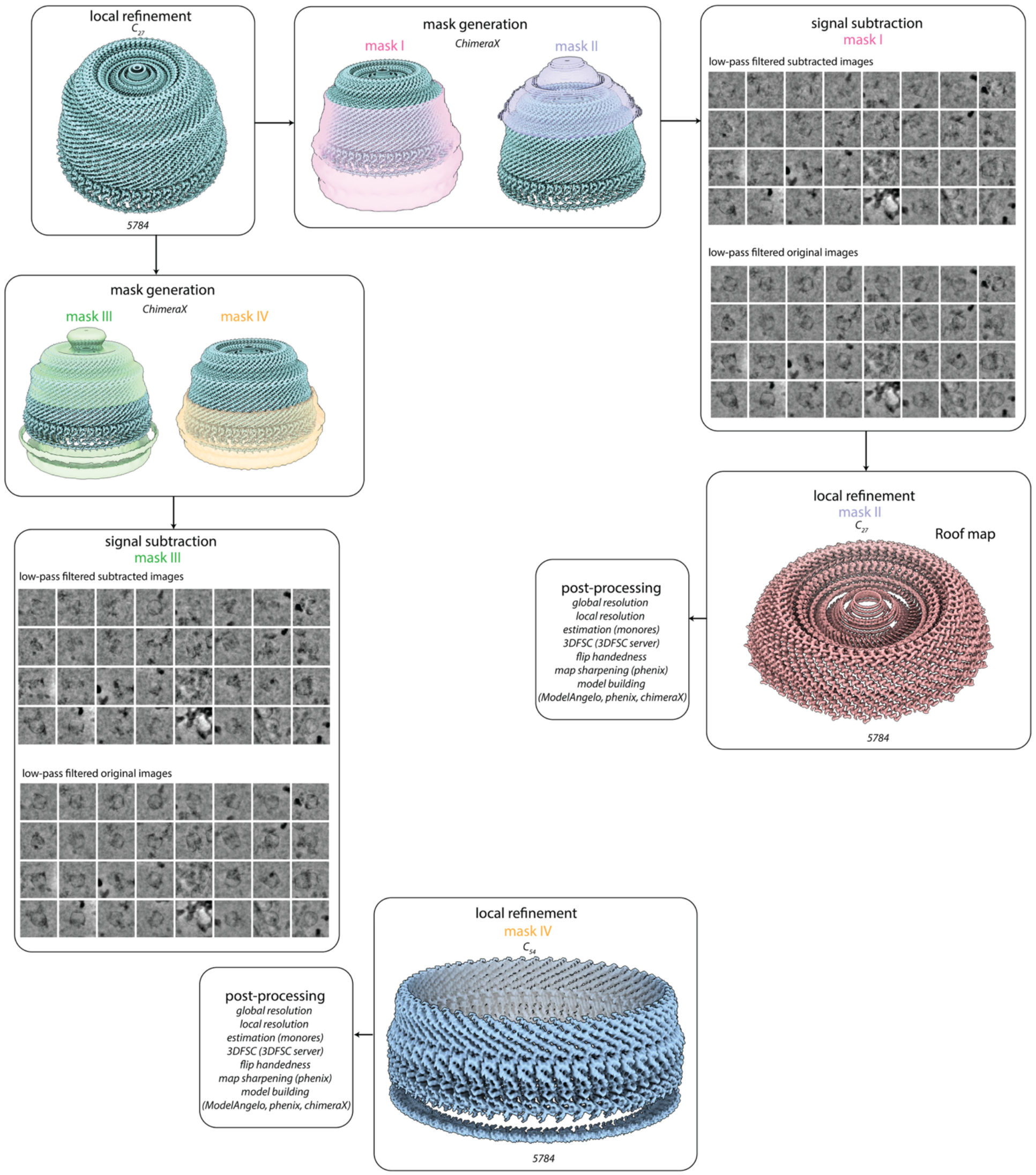
Signal subtraction and focused refinement of the YqiK roof and SPFH belt. Starting from the full-cage reconstruction, masks were generated in ChimeraX for signal subtraction and local refinement. For the roof, signal subtraction using mask I was followed by local refinement with mask II and C_27_ symmetry. For the SPFH belt, signal subtraction using mask III was followed by local refinement with mask IV and C_54_ symmetry. Representative low-pass-filtered particle images before and after subtraction are shown for each branch.

**Supplementary Figure 5.**
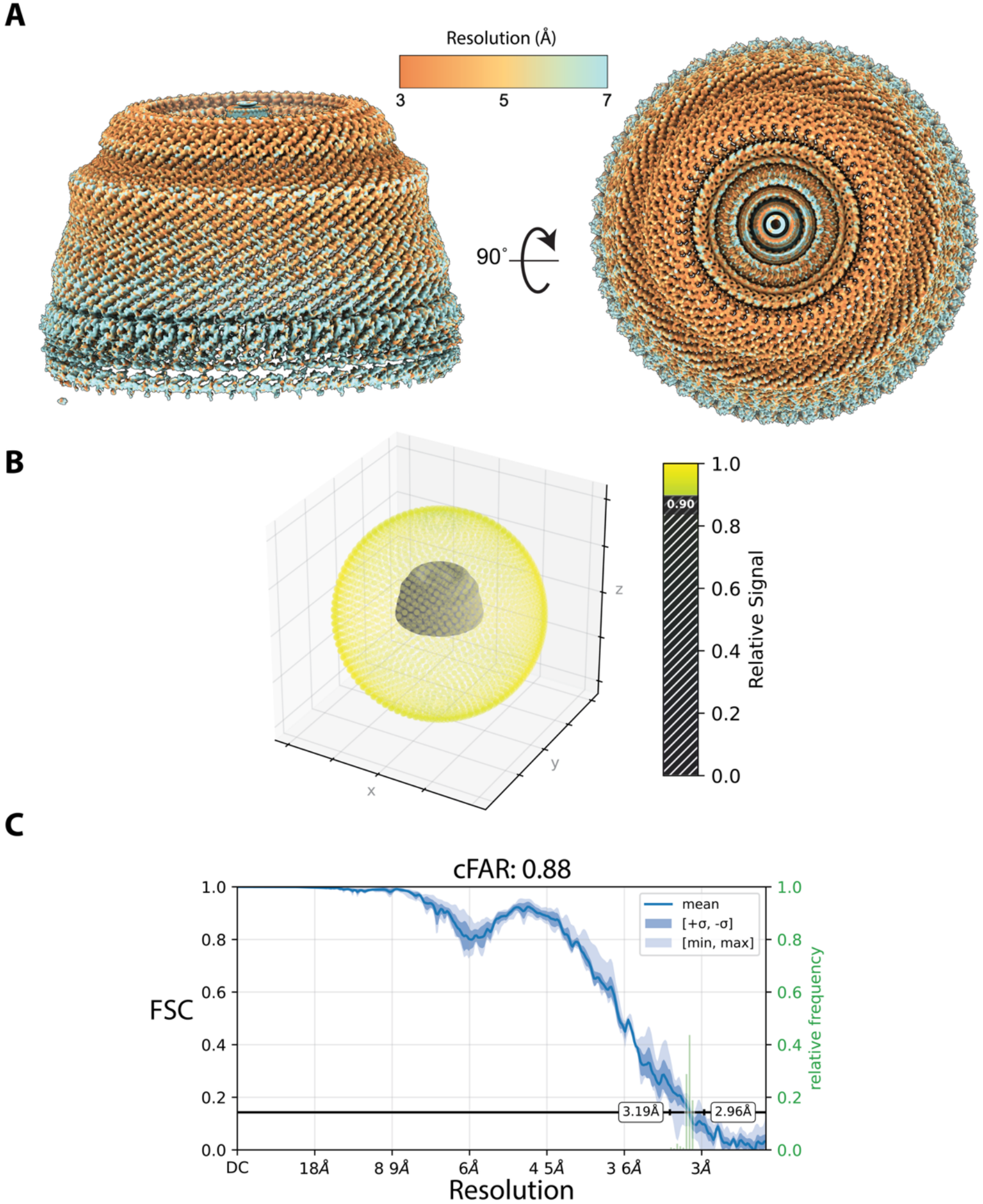
Local resolution and directional FSC analysis of the YqiK reconstruction. (**A**) Side and axial views of the cryo-EM map, related by a 90° rotation and colored by local resolution. (**B**) Three-dimensional distribution of relative signal across directions, colored according to the indicated scale. (**C**) Directional Fourier shell correlation (FSC) analysis. The blue curve shows the mean FSC; dark and light shaded regions indicate ±1 standard deviation and the minimum–maximum range, respectively. The horizontal black line marks the FSC threshold of 0.143. Green bars show the relative frequency of directional resolution estimates, with annotated limits of 2.96–3.19 Å. The reported cFAR value is 0.88.

**Supplementary Figure 6.**
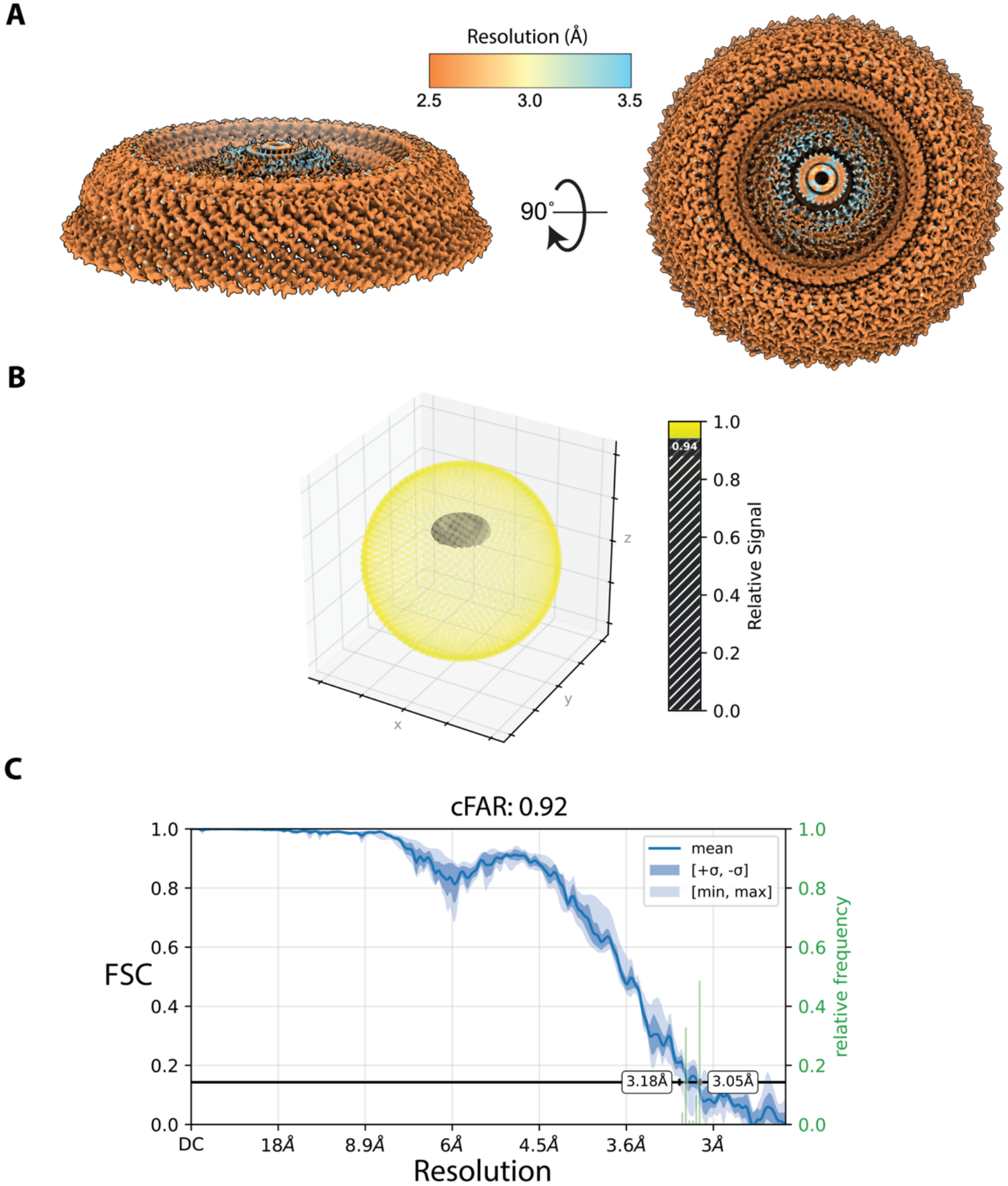
Local resolution and directional FSC analysis of the focused YqiK roof reconstruction. (**A**) Side and axial views of the focused roof map, colored by local resolution according to the indicated scale (2.5–3.5 Å). (**B**) Three-dimensional distribution of relative signal across directions, colored according to the indicated scale. (**C**) Directional Fourier shell correlation (FSC) analysis. The blue curve shows the mean FSC; dark and light shaded regions indicate ±1 standard deviation and the minimum–maximum range, respectively. The horizontal black line marks the FSC threshold of 0.143. Green bars show the relative frequency of directional resolution estimates, with annotated limits of 3.05–3.18 Å. The reported cFAR value is 0.92.

**Supplementary Figure 7.**
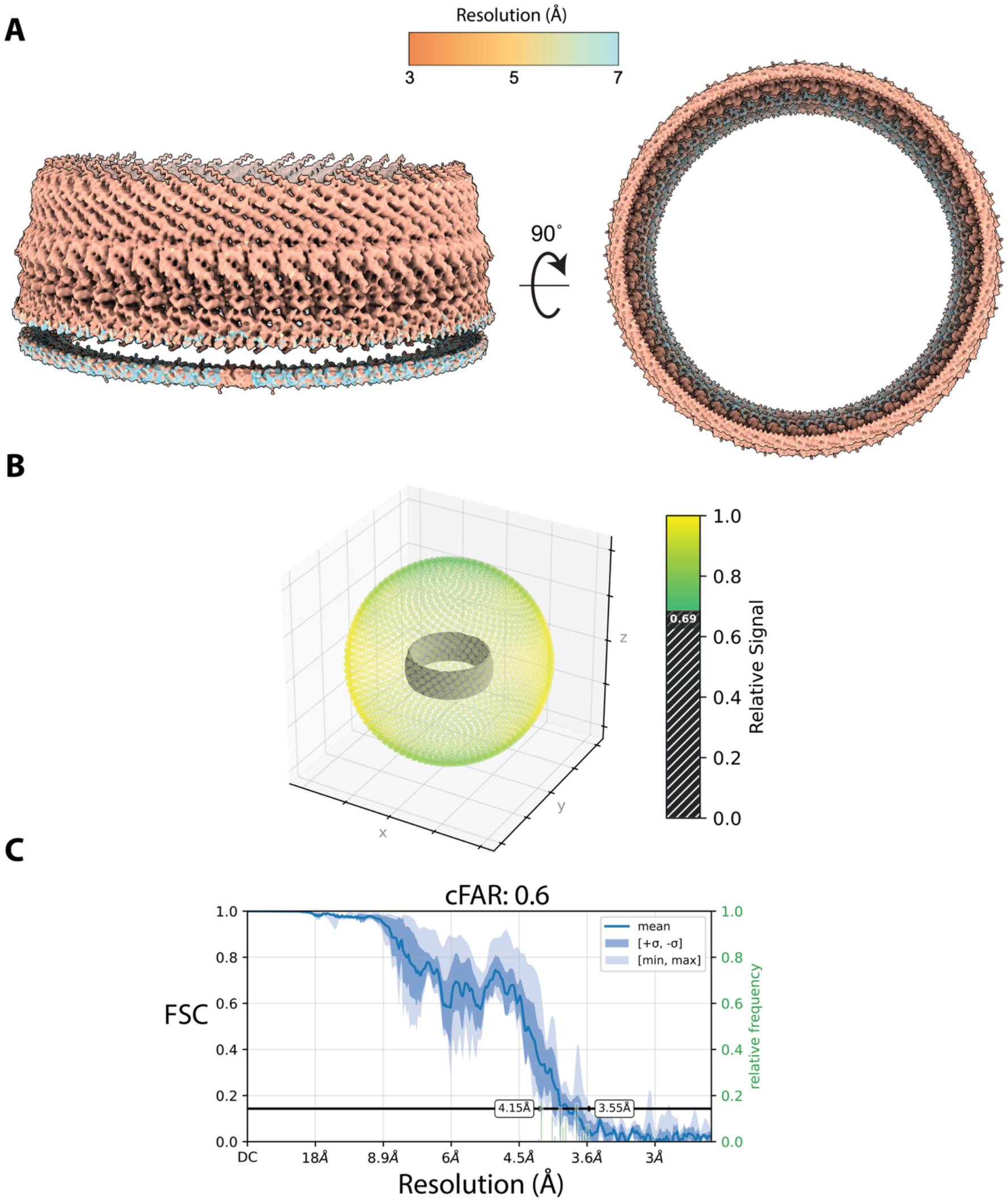
Local resolution and directional FSC analysis of the focused YqiK SPFH-belt reconstruction. (**A**) Side and axial views of the focused map, related by a 90° rotation and colored by local resolution according to the indicated scale (3–7 Å). (**B**) Three-dimensional distribution of relative signal across directions, colored according to the indicated scale. (**C**) Directional Fourier shell correlation (FSC) analysis. The blue curve shows the mean FSC; dark and light shaded regions indicate ±1 standard deviation and the minimum–maximum range, respectively. The horizontal black line marks the FSC threshold of 0.143. Green bars show the relative frequency of directional resolution estimates, with annotated limits of 3.55– 4.15 Å. The reported cFAR value is 0.60.

**Supplementary Figure 8.**
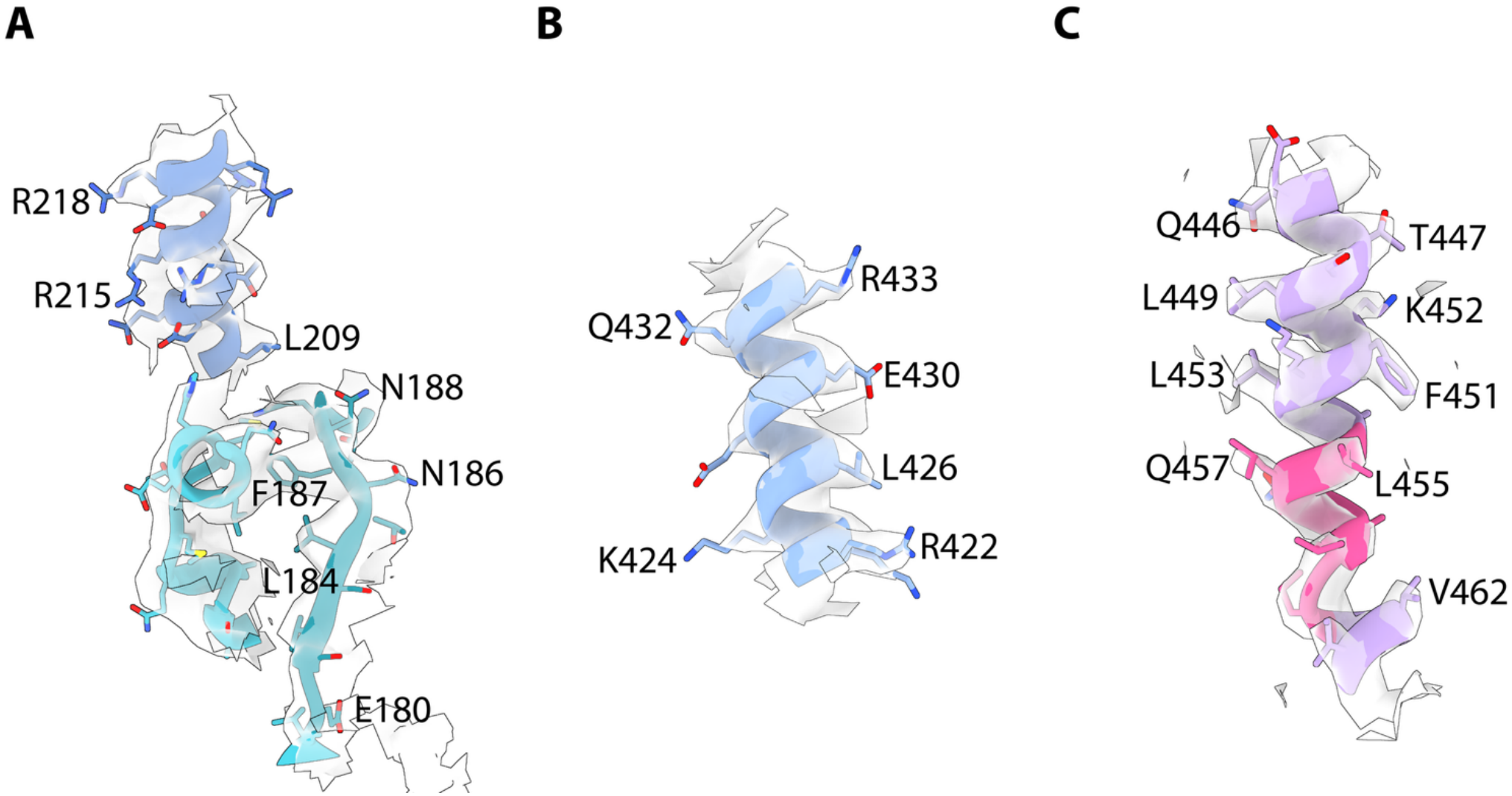
Representative cryo-EM density and model fit for YqiK. (**A**) SPFH domain and the base of the wall helix. (**B**) Upper wall helix. (**C**) Roof helix. Cryo-EM density is shown as a semitransparent grey surface, with the fitted atomic model shown as ribbons and selected side chains as sticks. Selected residues are labelled; nitrogen, oxygen and sulfur atoms are coloured blue, red and yellow, respectively.

**Supplementary Figure 9.**
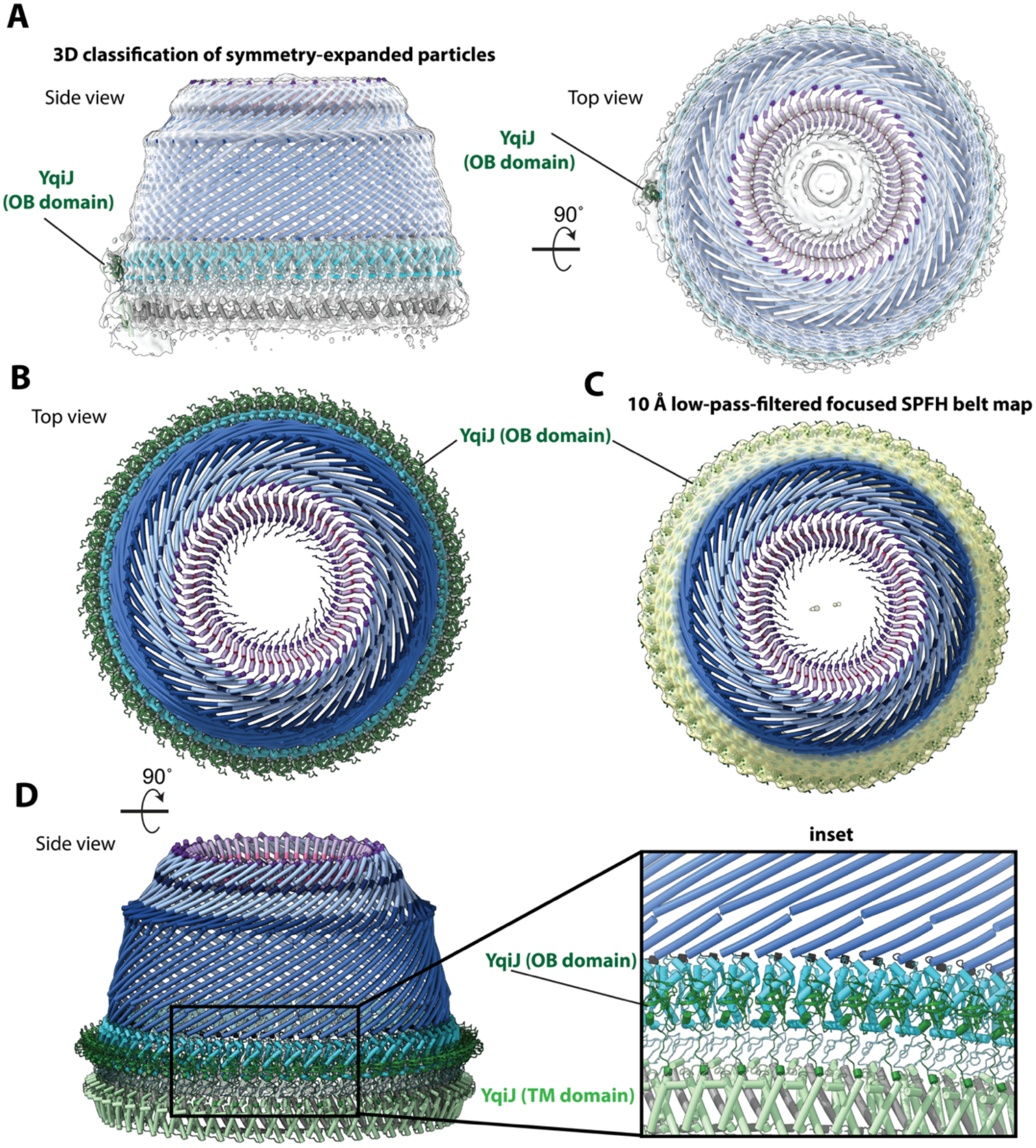
Cryo-EM density consistent with the predicted YqiJ-binding site on the YqiK cage. **(A)** Side and top views of a reconstruction obtained by focused 3D classification of symmetry-expanded particles (Fig. S3), shown as a semitransparent grey surface with the YqiK model and AlphaFold3-predicted YqiJ model superposed. Additional density on the outer face of the SPFH belt accommodates the predicted YqiJ OB-fold domain. (**B**) Top view of a model with one YqiJ per YqiK, illustrating the predicted arrangement of YqiJ OB-fold domains around the SPFH belt. (**C**) The focused SPFH-belt map, low-pass filtered to 10 Å and shown in yellow, superposed on the model in (**B**). Peripheral density overlaps the predicted positions of the YqiJ OB-fold domains. (**D**) Side view and enlarged inset showing the predicted placement of the YqiJ OB-fold domains (dark green) against the YqiK SPFH domains (cyan), with the YqiJ transmembrane domains (light green) alongside the YqiK transmembrane helices (grey).

**Supplementary Figure 10.**
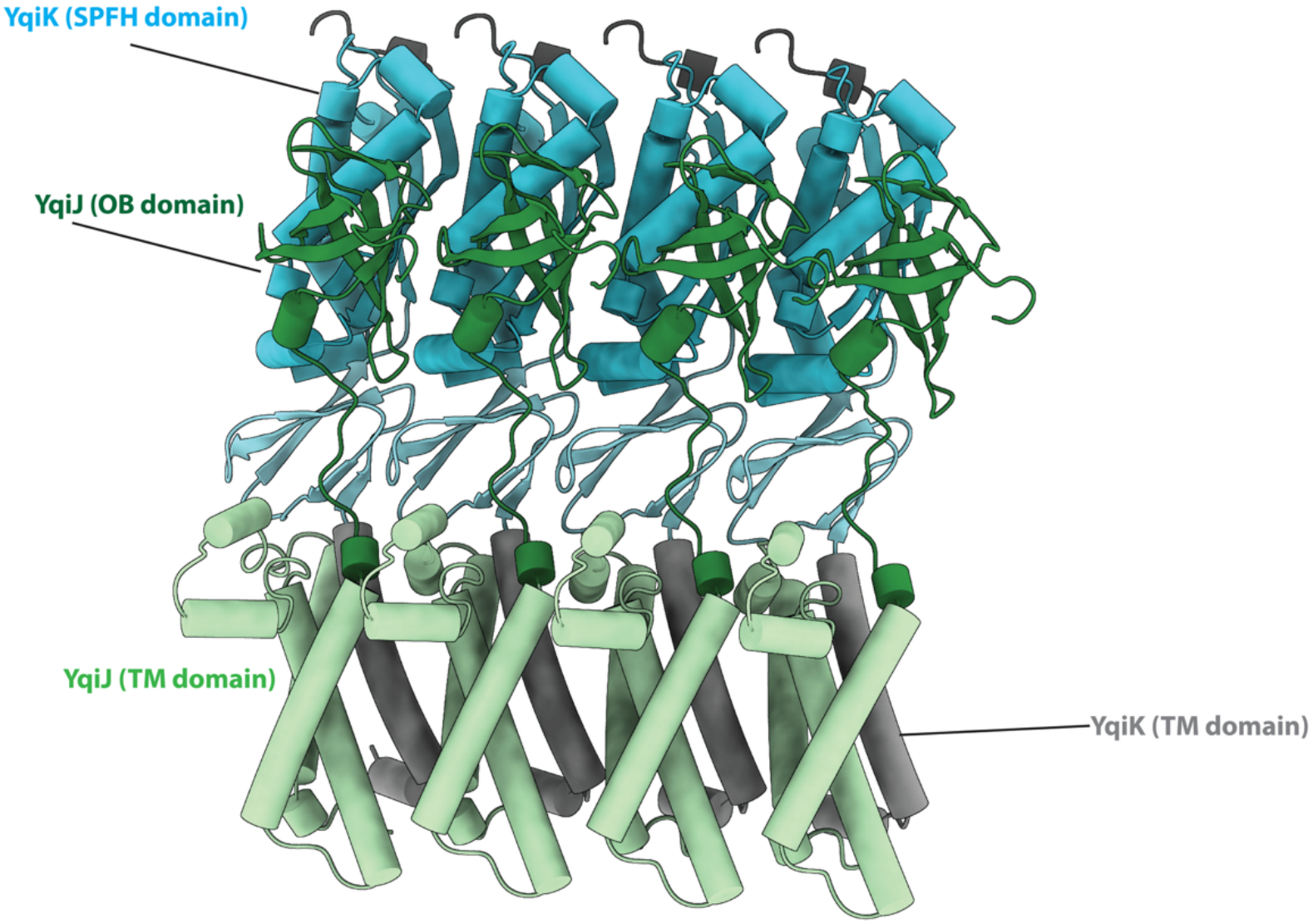
AlphaFold3-predicted arrangement of YqiK and YqiJ. Four adjacent YqiK–YqiJ units are shown in a model with one YqiJ per YqiK. The YqiJ OB-fold domain (dark green) lies against the outer face of the YqiK SPFH domain (cyan), while the YqiJ transmembrane domain (light green) lies alongside the YqiK transmembrane helix (grey).

**Supplementary Figure 11.**
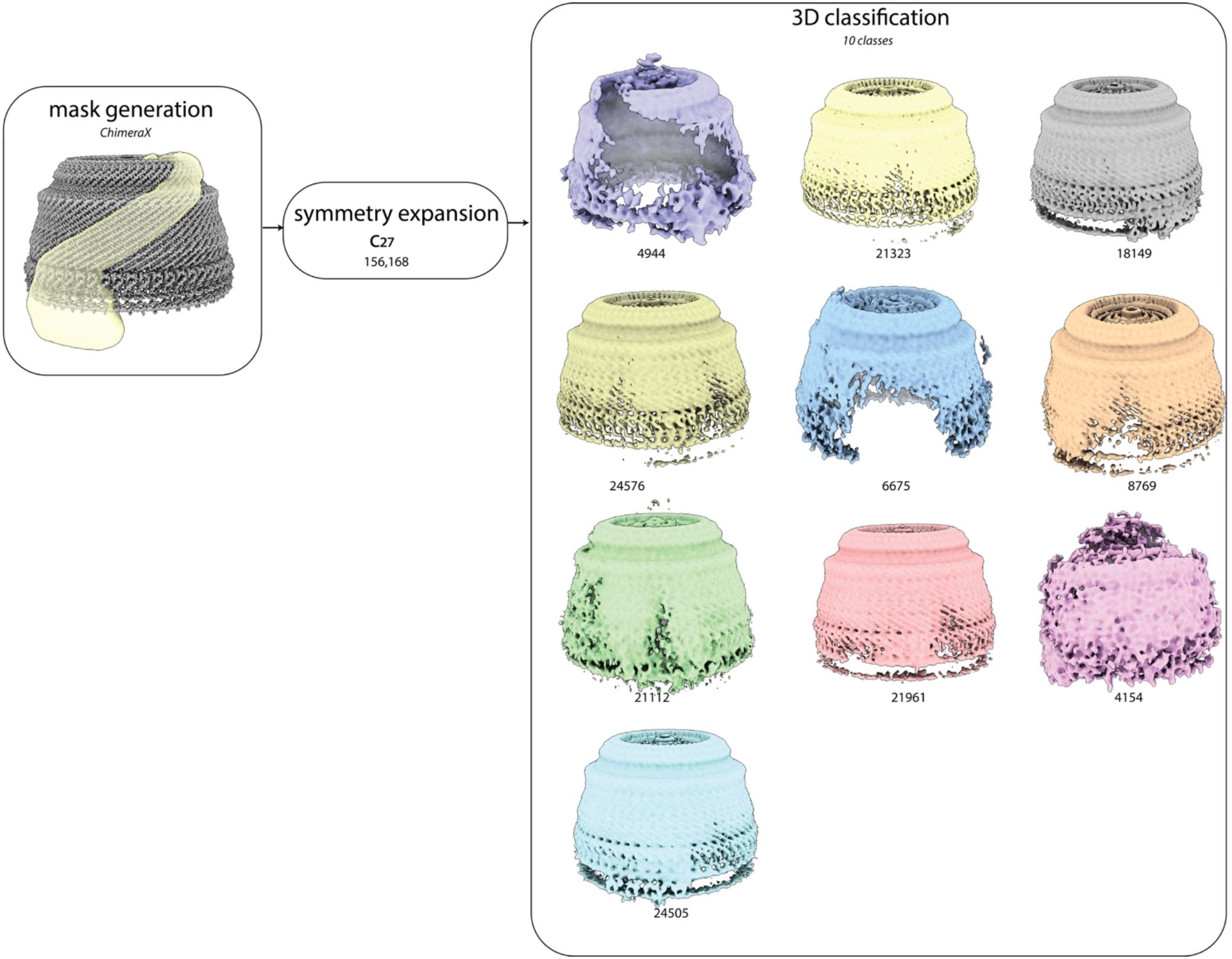
Focused 3D classification to identify potential open conformations of the YqiK cage. A mask encompassing a segment of the cage wall was generated in ChimeraX (yellow). C_27_ symmetry expansion of 5,784 particles yielded 156,168 symmetry-related particle copies, which were subjected to focused 3D classification into ten classes. The resulting reconstructions reveal variation in cage continuity, including classes with interrupted or missing wall density consistent with potential open or locally disordered conformations. Numbers beneath the maps indicate symmetry-expanded particle assignments, rather than counts of unique particles.

**Table S1.** Cryo-EM data collection, refinement and validation statistics.

|  | **Composite map + model** | **Consensus map** | **SPFH-focused map** | **Roof-focused map** |
|---|---|---|---|---|
| Accession codes | EMD-79940; PDB 40AZ | EMD-79930 | EMD-79931 | EMD-79939 |
| **Data collection and processing** |  |  |  |  |
| Microscope | Titan Krios G4 | Titan Krios G4 | Titan Krios G4 | Titan Krios G4 |
| Acquisition software | EPU | EPU | EPU | EPU |
| Voltage (kV) | 300 | 300 | 300 | 300 |
| Magnification | 130,000× | 130,000× | 130,000× | 130,000× |
| Pixel size (Å), collected / final map | 0.93 / 1.3053 | 0.93 / 1.3053 | 0.93 / 1.3053 | 0.93 / 1.3053 |
| Electron exposure (e<sup>−</sup>/Å<sup>2</sup>) | 50.12 | 50.12 | 50.12 | 50.12 |
| Defocus range (µm) | −0.4 to −2.2 | −0.4 to −2.2 | −0.4 to −2.2 | −0.4 to −2.2 |
| Movies (no.) | 18,457 | 18,457 | 18,457 | 18,457 |
| Initial particle images (no.) | 13,349 | 13,349 | 13,349 | 13,349 |
| Final particle images (no.) | 5,784 | 5,784 | 5,784 (signal-subtracted) | 5,784 (signal-subtracted) |
| Symmetry | — (composite) | C<sub>27</sub> | C<sub>54</sub> | C<sub>27</sub> |
| Map resolution (Å), FSC 0.143 | — (composite) | 3.06 (tightened mask); 3.16 (standard mask) | 3.69 (tightened mask); 3.88 (standard mask) | 2.96 (tightened mask); 3.00 (standard mask) |
| Unmasked resolution (Å), FSC 0.143 | — (composite) | 7.4 | 8.0 | 7.5 |
| **Refinement** |  |  |  |  |
| Map–model FSC 0/0.143/0.5 (Å, masked) | 2.9/3.0/3.6 | — | — | — |
| Correlation coefficient (mask/box/peaks/volume) | 0.81/0.82/0.72/0.79 | — | — | — |
| Chains / protein residues / non-hydrogen atoms | 54 / 25,731 / 201,393 | — | — | — |
| Mean B factor (Å<sup>2</sup>), protein (min/max) | 124.0 (42.9/421.6) | — | — | — |
| R.m.s. deviations, bonds (Å) / angles (°) | 0.003 / 0.522 | — | — | — |
| **Validation** |  |  |  |  |
| MolProbity score | 2.00 | — | — | — |
| Clashscore | 6.17 | — | — | — |
| Rotamer outliers (%) | 2.39 | — | — | — |
| Cβ outliers (%) | 0.00 | — | — | — |
| Ramachandran favored / allowed / disallowed (%) | 98.21 / 1.69 / 0.11 | — | — | — |

**Movie S1. Overall structure of the YqiK–YqiJ cage**. The 54-subunit atomic model is shown first, rotated about the cage axis and viewed down the axis, followed by a single subunit colored by region against the remaining 53 in grey: transmembrane helix, SPFH domain, lower and upper wall helix, bends 1 and 2, and the kink and roof. The composite cryo-EM map is shown last, colored by the fitted atomic model.

**Dataset S1. Spectronaut candidates report for FLAG-YqiJ eluates versus empty-vector controls**. Average log_2_ ratio, p-value and other information for each of the protein groups quantified in at least two replicates of both conditions. Related to Fig. 4A.

## Notes

### Competing Interest Statement

The authors have declared no competing interest.

